# Chloroquine and hydroxychloroquine as ACE2 blockers to inhibit viropexis of 2019-nCoV Spike pseudotyped virus

**DOI:** 10.1101/2020.06.22.164665

**Authors:** Nan Wang, Shengli Han, Rui Liu, Liesu Meng, Huaizhen He, Yongjing Zhang, Cheng Wang, Yanni Lv, Jue Wang, Xiaowei Li, Yuanyuan Ding, Jia Fu, Yajing Hou, Wen Lu, Weina Ma, Yingzhuan Zhan, Bingling Dai, Jie Zhang, Xiaoyan Pan, Shiling Hu, Jiapan Gao, Qianqian Jia, Liyang Zhang, Shuai Ge, Saisai Wang, Peida Liang, Tian Hu, Jiayu Lu, Xiangjun Wang, Huaxin Zhou, Wenjing Ta, Yuejin Wang, Shemin Lu, Langchong He

## Abstract

**Background:** The novel coronavirus disease (2019-nCoV) has been affecting global health since the end of 2019 and there is no sign that the epidemic is abating. The major issue for controlling the infectious is lacking efficient prevention and therapeutic approaches. Chloroquine (CQ) and Hydroxychloroquine (HCQ) have been reported to treat the disease, but the underlying mechanism remains controversial.

**Purpose:** The objective of this study is to investigate whether CQ and HCQ could be ACE2 blockers and used to inhibit 2019-nCoV virus infection.

**Methods:** In our study, we used CCK-8 staining, flow cytometry and immunofluorescent staining to evaluate the toxicity and autophagy of CQ and HCQ, respectively, on ACE2 high-expressing HEK293T cells (ACE2^h^ cells). We further analyzed the binding character of CQ and HCQ to ACE2 by molecular docking and surface plasmon resonance (SPR) assays, 2019-nCoV spike pseudotyped virus was also used to observe the viropexis effect of CQ and HCQ in ACE2^h^ cells.

**Results:** Results showed that HCQ is slightly more toxic to ACE2^h^ cells than CQ. Both CQ and HCQ could bind to ACE2 with *K_D_* =(7.31±0.62)e^−7^ M and (4.82±0.87)e^−7^ M, respectively. They exhibit equivalent suppression effect for the entrance of 2019-nCoV spike pseudotyped virus into ACE2^h^ cells.

**Conclusions:** CQ and HCQ both inhibit the entrance 2019-nCoV into cells by blocking the binding of the virus with ACE2. Our findings provide novel insights into the molecular mechanism of CQ and HCQ treatment effect on virus infection.

## Introduction

Chloroquine (CQ) and hydroxychloroquine (HCQ) are effective antimalarial drugs (White, 1996). The sole difference between their chemical structures is the presence of a hydroxymethyl group on HCQ as against a methyl group on CQ. The hydroxymethyl group enables HCQ to be absorbed in the human gastrointestinal tract faster and distributed in the body to larger extent than CQ (Rainsford et al., 2015; Schrezenmeier and Dorner, 2020). Since 2004, reports on the anti-viruses effects of CQ and HCQ have gradually increased. For example, CQ can inhibit the replication of SARS and HIV viruses *in vitro*, and it also has significant inhibitory effects on Borna, avian leukemia and Zika viruses. (Al-Bari, 2017; Keyaerts et al., 2004; Savarino et al., 2006). Therefore, CQ and HCQ have been considered broad-spectrum antiviral drugs.

Since the outbreak of 2019-nCoV (also called SARS-CoV-2) in 2020, there have been reports of CQ and HCQ used in clinical treatment. For example, clinical trial reports issued by 10 hospitals in China indicate that CQ may shorten the duration of the disease (Gautret et al., 2020). A small nonrandom clinical trial in France showed that HCQ combined with azithromycin has a significant therapeutic effect (Gautret et al., 2020), and it has been reported that CQ can effectively inhibit the deterioration of new coronary pneumonia, improve lung imaging performance, promote viral reversion and shorten the time of disease onset(Gao et al., 2020). It has been hypothesized that HCQ aerosols can be inhaled early in infection, allowing for the sufficient therapeutic effects on alveolar epithelial cells while avoiding the adverse effects of large oral doses(Klimke et al., 2020). However, it has also been reported that the combination of HCQ and azithromycin in 11 patients with severe 2019-nCoV infection has not achieved a positive clinical effect (Molina et al., 2020). Another observational study showed that HCQ had no effect on the intubation or composite endpoint of death (Geleris et al., 2020). There is still a lack of randomized controlled trials of HCQ in the treatment of patients with 2019-nCoV.

At present, it is generally believed that the 2019-nCoV enters the host cell by binding to ACE2 on the plasma membrane of the cells, causing infection (Hoffmann et al., 2020; Wrapp et al., 2020; Zheng et al., 2020). Therefore, blocking or antagonizing the ACE2 signaling pathway in susceptible cells should be beneficial in the prevention of 2019-nCoV infection (Wu et al., 2020). Abdelli *et al*. conducted a molecular docking experiment between CQ and ACE2 and found that CQ binds to ACE2 with low binding energy and forms a stable complex system (Abdelli et al., 2020). Studies have shown that ACE2 is a type I membrane-bound glycoprotein composed of 805 amino acids, mainly distributed in vascular endothelial cells, alveolar and renal tubular epithelial cells, and profoundly expressed in tissues such as heart, kidney, retina, and gastrointestinal tissue (Xiao et al., 2020). Flow cytometry and immunoprecipitation studies have shown that during alveolar epithelial cell infection with SARS virus, CQ and HCQ can prevent the binding of viral S protein to ACE2 by disrupting ACE2 terminal glycosylation (Brufsky, 2020; Vincent et al., 2005). Virus infection experiments *in vitro* confirmed that CQ could reduce the infection of cells by 2019-nCoV, and play a role in both the entry and post-entry stages of viral infection. At the same time, HCQ can effectively reduce the 2019-nCoV copy number (Wang et al., 2020b).

Recently, there have also been reports of adverse reactions of HCQ and CQ. A clinical trial of 90 patients with 2019-nCoV infection in the United States showed that 2019-nCoV positive patients receiving HCQ treatment had a higher risk of prolonged QTc, suggesting a risk of cardiotoxicity (Mercuro et al., 2020). At the same time, in a clinical trial of 197 2019-nCoV positive patients in China, CQ showed a significant therapeutic effect without severe adverse reactions (Mingxing et al., 2020). The above evidence suggests that the adverse effects of CQ treatment in 2019-nCoV posotive patients may be lower than that of HCQ. The curative effect and mechanism of the anti-2019-nCoV of CQ and HCQ are still controversial.

In this study, we found that CQ and HCQ can antagonize ACE2 and inhibit the entry of 2019-nCoV spike pseudotyped virus into ACE2 expressed HEK293T cells (ACE2^h^ cells).

## Materials and Methods

### Materials and Reagents

CQ, the purity of 98%, was from Macklin (Shanghai, China), HCQ, the purity of 98 %, was provided by Energy Chemical, (Shanghai, China). Dulbecco’s Modification of Eagle’s Medium (DMEM) with high glucose (Cat. No. SH30022.01), and fetal bovine serum (Cat. No. 16140071) were from HyClone (Logan, UT, USA). Penicillin–streptomycin solution was obtained from Xi’an Hat Biotechnology Co., Ltd (Xi’an, China). Protease inhibitor and phosphatase inhibitor cocktails were purchased from Roche Diagnostic (Mannheim, Germany). The 5×loading buffer was purchased from Thermo Fisher Scientific, lnc. (MA, USA), and SDS-PAGE was from Pioneer Biotech Co., Ltd (Xi’an, China). Polyvinylidene fluoride membranes were from Hangzhou Microna Membrane Technology Co., Ltd (Hangzhou, China). Tween-20 was provided by Shaanxi Pioneer Biotech Co., Ltd (Xi’an, China). Enhanced Chemiluminescence (ECL) kit was from Proteintech Group, lnc (Rosemont, USA). Annexin V-FITC/PI Apoptosis Detection Kit (Cat. No. A005-3) and Cell Counting Kit were purchased from 7Sea Pharmatech Co., Ltd (Shanghai, China), the 2019-nCoV spike pseudotyped virus (Cat: PSV001) was purchased from Sino Biological (Beijing, China)

### Cell culture

HEK293T cells, human airway epithelial cells (HSAEpC), alveolar type II epithelial cells (AT2), and eosinophilic leukemia (EOL-1) cells were from ATCC. ACE2^h^ cells were constructed by Genomeditech (Shanghai, China). HSAEpC and AT2 cells were maintained in DMEM with high glucose containing 10% FBS and 1% penicillin-streptomycin; EOL-1 cells were kept in 1640 medium containing 10% FBS and 1% penicillin-streptomycin; ACE2^h^ cells were maintained in DMEM with high glucose medium containing 10% FBS, 1% penicillin-streptomycin, and 4 μg/mL puromycin and cultured at 37°C in a 5% CO^2^ incubator.

### Cytotoxicity assay

Cell viability was determined following the manufacturer’s instructions. Briefly, ACE2^h^ cells were seeded into 96-well plates at a density of 5 × 10^3^ cells per well and then treated with different concentrations of CQ or HCQ (0, 0.1, 1, 10, 50, 100, 200, 300 and 400 μM) for 24 h, then 10 μL of Cell Counting Kit solution was added to each well followed by 2 h of incubation. Relative cell viability was assessed by measuring the absorbance at 450 nm using a microplate reader (Bio-Rad, Carlsbad, CA, USA). The survival rate of ACE2^h^ cells was calculated using the following formula:

[(OD_Treated_ − OD_Blank_) / (OD_Control_ − OD_Blank_)] × 100%. The time-dependent effects (6, 12, 24 and 48 h) of HCQ and CQ on ACE2^hi^ cell viability at low concentrations (10 and 20 μM) were also observed using the same method.

### Apoptosis assay

ACE2^h^ cells were seeded in a six-well plate and treated with different concentrations of CQ and HCQ (0, 10, 20 and 40 μM) for 24 h. Cells were collected and washed with PBS and resuspended in 400 μL of 1 × binding Buffer. Annexin V-FITC (5 μL) was added to the cells and incubated 26 °C in the dark for 15 min. PI (10 μL) was added to the cells and incubated in an ice bath for 5 min. Detection was performed within 30 min. The excitation wavelength of the flow cytometer (Accuri C6 Plus, BD Biosciences, Beijing, China) was 488 nm, and the emission wavelength was 530 nm to detect FITC, while PI was detected at 575 nm. Normal cells had low fluorescence intensity. Apoptotic cells had strong green fluorescence, and necrotic cells had double staining with green and red fluorescence.

### Western blotting

Total proteins from different cells were extracted in ice-cold conditions using RIPA lysis buffer containing 10% protease inhibitor and a phosphatase inhibitor cocktail. The protein concentration was determined using a BCA Protein Quantification kit according to the manufacturer’s instructions. The protein in the cell lysates was denatured by boiling the samples for 5 min with a 5 × loading sample buffer and equal amounts of protein were separated on a 10% gel using SDS-PAGE. The separated proteins were transferred onto polyvinylidene fluoride membranes and blocked by constant stirring with 5% nonfat milk in Tris-buffered saline containing Tween-20. The membranes were then incubated overnight at 4°C with the following primary antibodies: anti-ACE2 (1:500, EPR4435, Abcam), anti-LC3 (1:1000, #2775, Cell Signaling Technology [CST]) and anti-GAPDH (1:2000, a#2118, CST). The membranes were washed three times with TBST and then incubated with secondary antibodies (a dilution of 1:20,000 in TBST) for 1 h at 37°C. The membranes were washed three times with TBST for 10 min and developed using ECLkit. A Lane 1 DTM transilluminator (Beijing Creation Science, Beijing, China) was used to capture the images of the developed blots, and Image-Pro Plus 5.1 software (Rockville, MD, USA) was used to quantify the protein levels.

### Immunofluorescence assays

ACE2^h^ cells (2×10^3^) were seeded on 24 mm×24 mm coverslips. and incubated overnight at 37 °C with 5 % CO_2_.10 μM, 20 μM or 40 μM CQ and HCQ were added to the slides and treated for 24 h. The slides were then fixed with 4 % paraformaldehyde, followed with 0.5% Triton X-100 for 5 min and 5% BSA solution for 1 h at 26°C after washing three times with PBS. The cells were then continuously incubated with LC3 primary antibody at 37°C for 3 h, and the fluorescent secondary antibody at 26°C for 2 h followed with TRITC-Phalloidin stain for 30 min at 26°C. Finally, the cells were mounted with 50 μL of DAPI-containing anti-fluorescence quenching reagent. All the cells were observed using a laser confocal fluorescence microscope.

### Docking Studies

Molecular docking studies were carried out using SYBYL-X 2.0 version. The small molecules and X-ray crystal structure of the protein (PDB code: 6M0J) were imported. Water molecules were removed and hydrogen was added. Tripos force field and Pullman charge were applied to minimize. CQ and HCQ were depicted by the Sybyl/Sketch module (Tripos Inc.), optimized by Powell’s method with the Tripos force field with convergence criterion at 0.005 kcal/(Å mol), and assigned using Gasteiger–Hückel method.

### Surface plasmon resonance assay

For assessment of surface plasmon resonance (SPR), ACE2 protein with a 6-his tag (30 μg/mL) was fixed on a carboxyl sensor chip (Nicoya, Canada) by capture-coupling. Then, CQ or HCQ at 6.25, 12.5, 25, 50 and 100 μM was injected sequentially into the chamber in PBS running buffer. The interaction of ACE2 with the fixed small molecules was detected using Open SPR^TM^ (Nicoya Lifesciences, Waterloo, Canada) at 25°C. The binding time and disassociation time were both 250 s, the flow rate was 20 μL/s, and the chip was regenerated with hydrochloric acid (pH 2.0). A one-to-one diffusion-corrected model was fitted to the wavelength shifts corresponding to the varied drug concentration. The data were retrieved and analyzed using TraceDrawer.

### Detection of 2019-nCoV spike pseudotyped virus entry into ACE2^h^ cells

For this process, 5 × 10^4^ of ACE2^h^ cells in 50 μL DMEM per well were seeded into white 96-well plates. The cells were cultured in a 37 °C incubator containing 5% CO_2_ for 2 h. Medium (25 μL) was aspirated carefully from 96 wells, 25 μL medium containing the corresponding dose of the medicine was added and incubated for 2 h. Then 5 μL of 2019-nCoV spike pseudotyped virus was added (Sino Biological, PSC001), and incubated in a 37 °C incubator containing 5% CO_2_ for 4 h followed with adding 100 μL of complemented DMEM per well. After 6-8 h of further infection, the culture medium containing the virus was removed and replaced by 200 μL of fresh DMEM, and incubated continuously at 37℃ for 48 h, the culture medium was aspirated and 20 μL of cell lysate was added from the Luciferase Assay System (Promega, E1500) to each well, Following this, 100 μL of luminescence solution was added to wells before the luciferase luminescence detection, chemiluminescence was detected by a microplate reader under 560 nm, with exposure time of 1 s.

### Statistical analysis

Data are presented as the mean ± standard error of the mean (SD) and were statistically analyzed using analysis of variance (ANOVA). Two-tailed tests were used for comparisons between two groups, and differences were considered statistically significant at *p* <0.05.

## Results

### Effect of CQ and HCQ on ACE2^h^ cell viability

The expression of ACE2 protein in human lung and bronchial-related cells was higher than that in HEK293T cells. The expression of ACE2 protein in ACE2^h^ cells was significantly higher than that in other cells, indicating that ACE2^h^ cells were successfully constructed. It has been reported that AT2 cells express the highest ACE2 receptors in lung and bronchial cells (Zou et al., 2020). We confirmed that the highest expression of the ACE2 protein occurred in AT2 cells. In addition, this is the first report that EOL-1 cells also express the ACE2 protein (Figure 1A).

**Figure 1.**
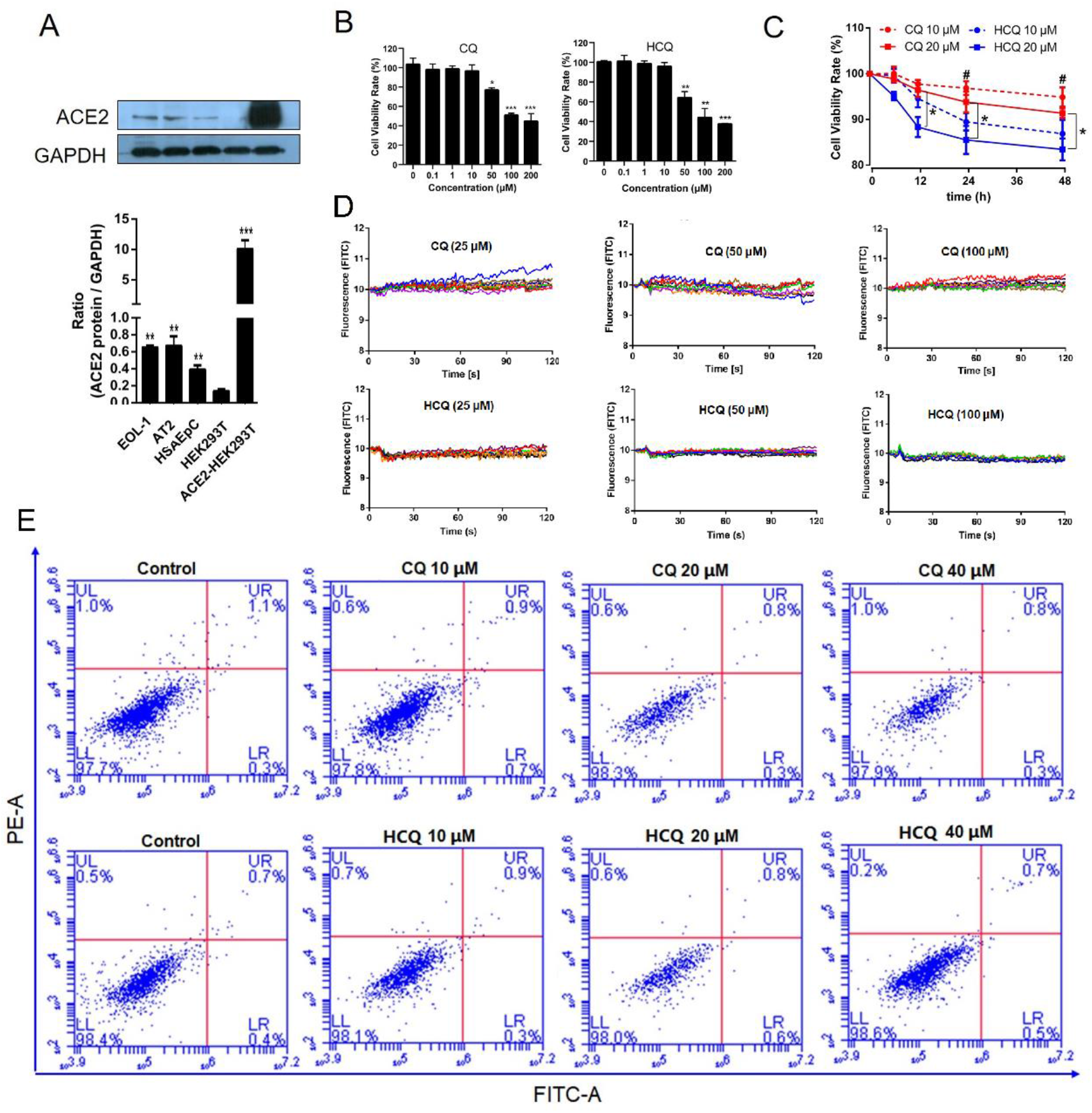
Effect of CQ and HCQ on the viability of ACE2^h^ cells. A. Western blotting analysis of the expression levels of ACE2 protein in EOL-1 cells, AT2 cells, HSAEpC cells, and ACE2^h^ cells. B. Viability of ACE2^h^ cells treated with CQ or HCQ for 24 h. C. The toxicity of HCQ and CQ on ACE2^h^ cells at different time points. D. Calcium (Ca2+) flux change in ACE2^h^ cells. E. The apoptosis of ACE2^h^ cells treated with CQ or HCQ for 24 h. The experiments were repeat three times. Data are presented as mean ± S.D. (\**p* < 0.05, \*\**p* < 0.01, \*\*\**p* < 0.001, compared with HEK293T, or concentration was 0, or HCQ 20 μM, ^#^*p* < 0.05 compared with HCQ 10 μM at corresponding time points).

As shown in Figure 1B, CQ and HCQ had no significant effect on the activity of ACE2^h^ cells when the concentration was less than 50 μM, and the survival rate of ACE2^h^ cells could be reduced in a dose-dependent manner when the concentration was above 50 μM. The inhibition of HCQ on the activity of ACE2^h^ cells was more significant than that of CQ. It can be concluded that the toxicity of HCQ was higher than that of CQ on ACE2^h^ cells at different time points at the same concentrations (Figure 1C). At a concentration of 20 μM, the statistical difference appeared at 6 h. Ca^2+^ is an essential second messenger in several cell pathways, as shown in Figure 1D, and CQ or HCQ rarely affects Ca^2+^ influx change in ACE2^h^ cells. Figure 1E shows that within 24 h, the concentrations of both drugs had no significant effect on apoptosis.

### CQ and HCQ induce LC3-mediated autophagy in ACE2^h^ cells

Autophagosome is a spherical structure and as an essential marker for autophagy, and LC3 is known to be stably associated with the autophagosome membranes. LC3 includes two forms LC3-I and LC3-II, LC3-I is found in the cytoplasm, whereas LC3-II is membrane-bound and converted from LC3-I to initiate formation and lengthening of the autophagosome. Therefore, to investigate the effects of CQ and HCQ induced autophagy on ACE2^h^ cells, FITC-LC3, TRITC-Phalloidin and DAPI staining were used. Activating lysosomal (green) and filamentous actin (F-actin, red) was detected after stimulation with 10, 20 and 40 μM of CQ and HCQ in ACE2 cells (Figure 2A).

**Figure 2.**
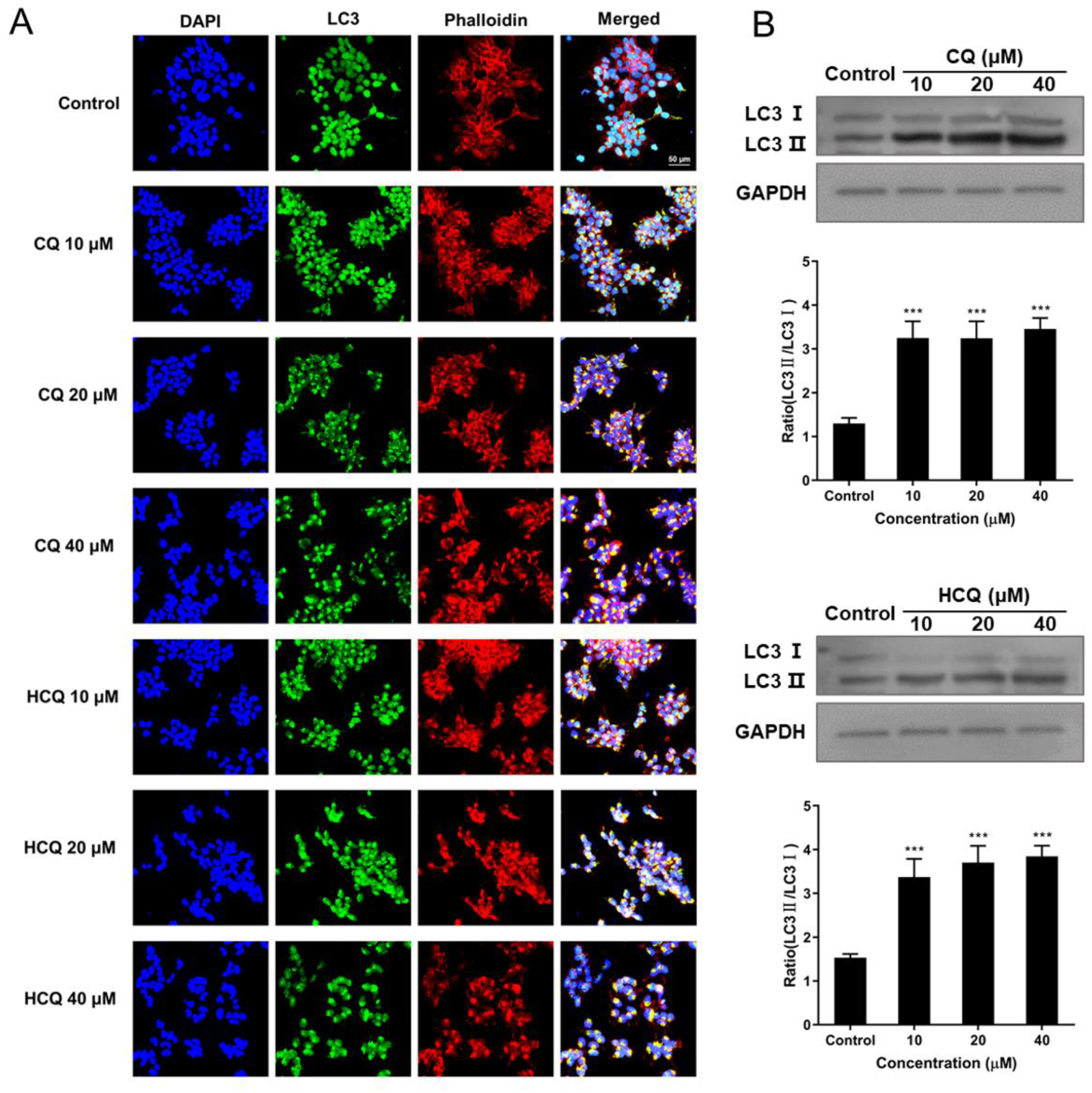
Effects of CQ and HCQ on the LC3 levels of ACE2^h^ cells. ACE^h^ cells were treated with different doses of CQ or HCQ for 24 h. (A) Effects of CQ and HCQ on the fluorescent staining of FITC-LC3 and TRITC-Phalloidin in ACE^h^ cells. (B) The representative blots of autophagy proteins and changes of LC3-II/LC3-I ratio. The experiments were repeat three times. Data are presented as mean ± S.D. \**p* < 0.05, \*\**p* < 0.01, \*\*\**p* < 0.001 compared with control.

We further pretreated ACE2^h^ cells with CQ and HCQ, and measured the expression of ACE2^h^ cells autophagy proteins LC3-I and LC3-II by Western blotting. We found that the expression level of LC3 and LC3-II increased in CQ and HCQ-treated ACE2^h^ cells (Figure 2B). The protein level of the LC3-II/LC3-I ratio was significantly increased compared to the control group (Figure 2B). All of these results suggested that CQ and HCQ could induce LC3-mediated autophagy in ACE2^h^ cells.

### Binding characteristics of CQ and HCQ with ACE2

The SARS-CoV-2 virus infects its host cells through binding to the ACE2 protein followed by cleavage of the spike protein by human TMPRSS2, we focused on whether CQ or HCQ could bind with ACE2. A virtual molecular docking test was performed to investigate the binding character of CQ and HCQ with ACE2. The chemical structure of both drugs are showed in Figure 3A. Figure 3B shows that both CQ and HCQ can bind to R393 and D350 (both in green) of ACE2 with their quinoline and imino groups. In addition, due to the replacement of a methyl group by a hydroxymethyl group, HCQ can form two additional hydrogen bonds with D350 and S44 (in red).We further used SPR to confirm the binding between CQ or HCQ and ACE2. The binding constant *K_D_* of these two compounds and ACE2 protein were (7.31±0.62)e-7 and (4.82±0.87)e-7 M respectively (Figure 3C).

**Figure 3.**
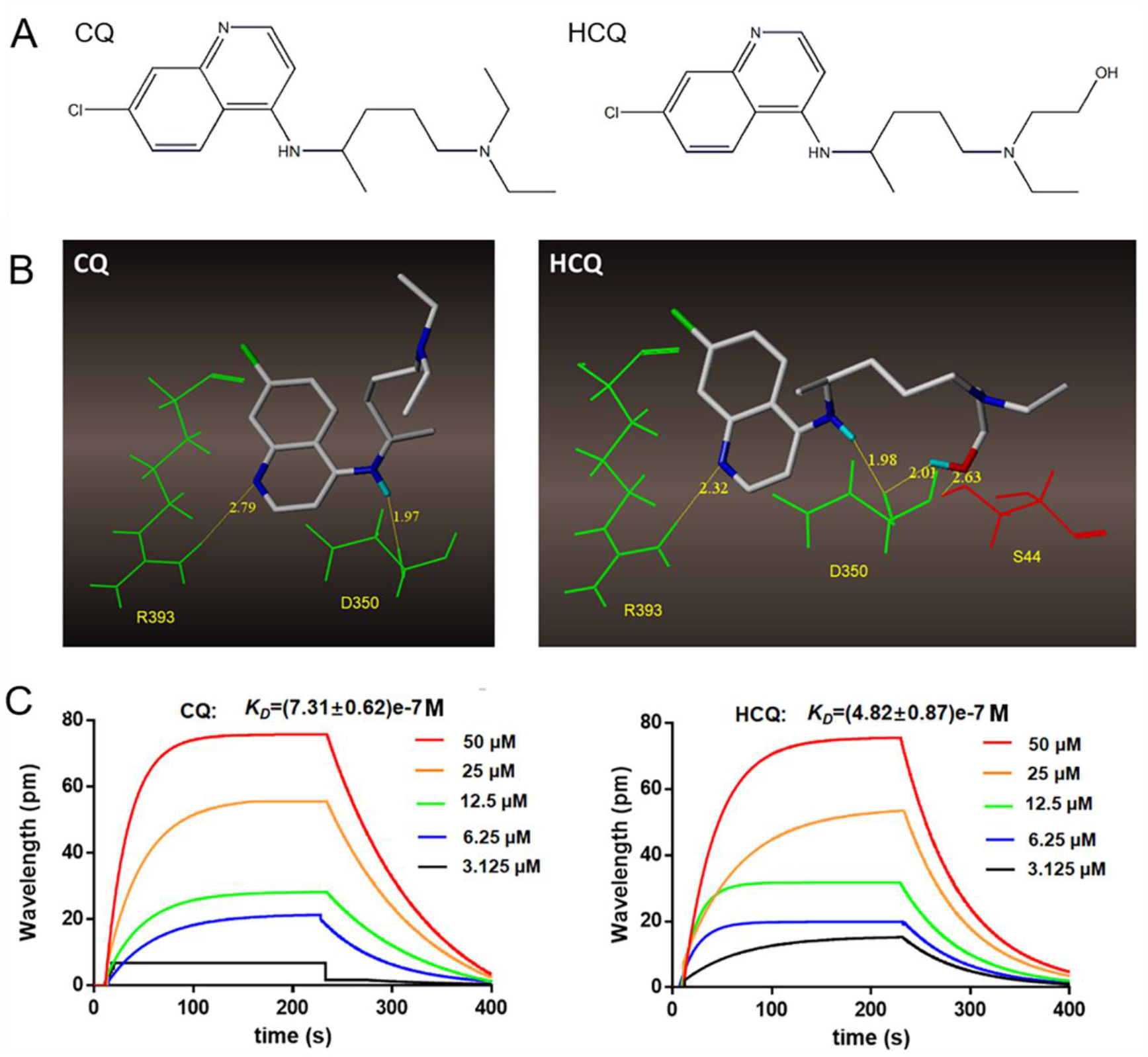
Binding character of CQ and HCQ with ACE2. A. Structural formulas of CQ and HCQ. B. SPR analysis of CQ or HCQ and ACE2. C. Molecular docking of CQ and HCQ with ACE2.

### CQ and HCQ suppressed the entrance of 2019-nCoV spike pseudotyped virus into ACE^h^ cells

ACE^h^ cells infected only with 2019-nCoV spike pseudotyped virus were considered as controls, and the luciferase luminescence value of the control was defined as 1. Under treatment of 0.625 μM, 1.25 μM, 2.5 μM, 5 μM, 10 μM, and 20 μM CQ, the 2019-nCoV spike pseudotypes virus entrance ratio were reduced to 86±0.11, 69±0.13, 62±0.19, 56±0.13, 44±0.18 and 23±0.10%, respectively, when treated by the same dosage of HCQ, the ratios were 77±0.07, 58±0.12, 53±0.09, 44±0.08, 35±0.05, and 29±0.05% respectively (Figure 4). The ability of the 2019-nCoV spike pseudotyped virus to enter ACE2^h^ cells was significantly reduced after treatment with both CQ and HCQ.

**Figure 4.**
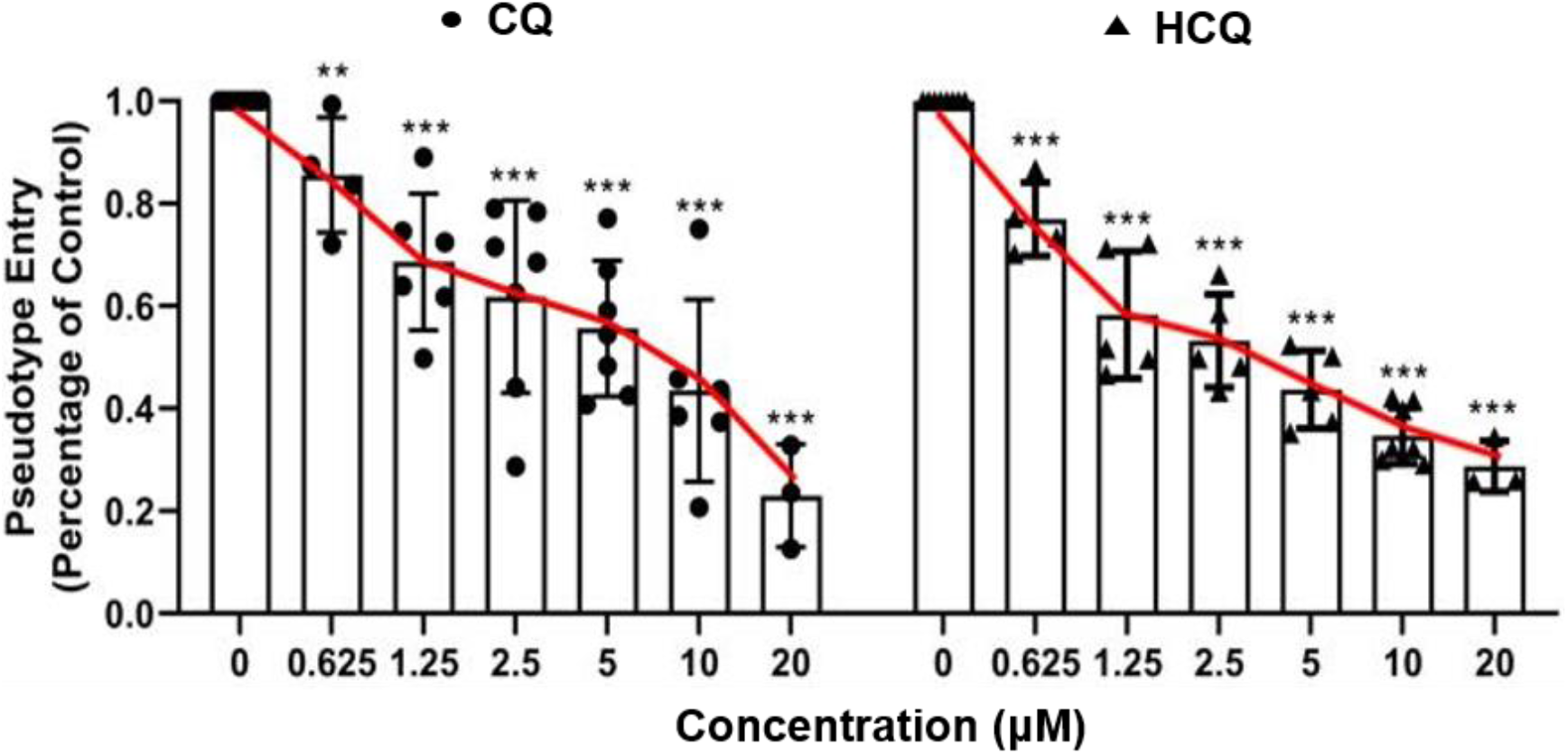
Effect of CQ and HCQ on the entrance of 2019-nCoV spike pseudotyped virus into ACE2^h^ cells. The experiments were repeat three times. Data are presented as mean ± S.D. \**p*<0.05, \*\**p*<0.01, \*\*\**p*<0.001 compared with group 0.

## Discussion

2019-nCoV is globally prevalent in 2020(Wang et al., 2020a), and there are currently no specific drugs against the virus(Lei et al., 2020). ACE2 is the target receptor of 2019-nCoV (Yan et al., 2020), and CQ and HCQ have shown certain efficacy in clinical use(Fantini et al., 2020; Ferner and Aronson, 2020; Meo et al., 2020). This study confirmed that both CQ and HCQ can interact with ACE2 and inhibit the entry of pseudoviruses.

CQ and HCQ have traditionally been used as anti-malaria drugs (White, 2007; White et al., 2014). They can easily induce a resistance to the malaria paraside (Gasquet et al., 1995), and are still recommnented for use solely against malaria. Further studies are needed to test the senstivity to local malaria strains since it is safe, efficient and cheap (Gutman et al., 2017). Recently, they have also been used commonly as immune modification drugs to treat autoimmune disorders (Plantone and Koudriavtseva, 2018). The underlying mechanism against malaria seems clear, but the mechanism of anti-inflammation is still under investigation. An increasing number of people believe that CQ and HCQ protect of lysosomes and could change pH values in lysosome (Mauthe et al., 2018). The two drugs have been reported to treat certain viral infections, but the antivirus mechanism remains unclear (Savarino, 2011). In Zika virus infection, CQ has been reported to be an endocytosis-blocking agent, and can inhibit the virus in different cell models (Delvecchio et al., 2016). Similarly, 2019-nCoV was driven by endocytosis after binding to ACE2.

CQ and HCQ possess structural differences, and the presence of the hydroxymethyl group in HCQ allows it to form additional hydrogen bonds with ACE2 according to the molecular docking results. These different modes may finally reveal different bioactivities and affinitis of CQ and HCQ on ACE2. Based on the above results, we further analyzed the binding strength of CQ and HCQ to the ACE2 protein, and found that both CQ and HCQ display strong binding to the ACE2 protein.

Virus entry into cells is a critical step in the process of virus infection (Shang et al., 2020). However, novel coronavirus research is greatly limited by the need to achieve a laboratory safety level of 3 or above for direct research using virus strains(Nie et al., 2020). A pseudovirus is a retrovirus that can integrate the membrane glycoproteins of a different kind of virus to form an external viral membrane, while retainings the genomic characteristics of the retrovirus itself. Construction of the 2019-nCoV pseudovirus that can only infect cells once, ensure safety and allows simulation the process of virus invasion into the cell to detect whether drugs have antiviral activity *in vitro* (Ou et al., 2020). Therefore, we use 2019-nCoV pseudovirus as an infection model to assess the antiviral effects of CQ and HCQ. We confirmed that both CQ and HCQ have the ability to suppress the entrance of 2019-nCoV spike pseudotypes virus into ACE2^h^ cells. 2019-nCoV uses ACE2 for cellular entry. Thus, CQ and HCQ could be good inhibitors to block 2019-nCoV infection of human cells expression ACE2. However, the difference in the inhibitory effect of these two drugs on 2019-nCoV needs further study.

Our study revealed that CQ and HCQ as ACE2 blockers inhibit the entrance of 2019-nCoV pseudovirus into the cells, providing new insights into the use of CQ and HCQ for 2019-nCoV treatment and further control.

## Author contributions

Nan Wang: Methodology, Supervision, Visualization, Writing - Original Draft; Shengli Han: Methodology, Supervision; Rui Liu: Investigation, Validation, Formal analysis; Liesu Meng: Formal analysis, Data Curation; Huaizhen He: Data Curation; Yongjing Zhang: Investigation, Visualization; Cheng Wang: Visualization, Software; Yanni Lv: Investigation, Validation; Jue Wang: Investigation, Visualization; Xiaowei Li: Investigation, Validation; Yuanyuan Ding: Investigation; Jia Fu: Investigation; Yajing Hou: Investigation; Wen Lu: Investigation; Weina Ma: Methodology; Yingzhuan Zhan: Data Curation; Bingling Dai: Methodology; Jie Zhang: Methodology; Xiaoyan Pan: Methodology; Shiling Hu: Investigation; Jiapan Gao: Investigation; Qianqian Jia: Investigation; Liyang Zhang: Investigation; Shuai Ge: Investigation; Saisai Wang: Investigation; Peida Liang: Investigation; Tian Hu: Investigation; Jiayu Lu: Investigation; Xiangjun Wang: Investigation; Huaxin Zhou: Investigation; Wenjing Ta: Investigation; Yuejin Wang: Investigation; Shemin Lu: Resources, Supervision, Data Curation, Formal analysis, Writing - Review & Editing; Langchong He: Resources, Supervision, Conceptualization, Funding acquisition

## Acknowledgement

This work was cofounded by National Natural Science Foundation of China (Grant number: 81930096) and Fundamental Research Funds for the Central Universities of China (Grant number: xzy032020042).

